# The intrinsically disordered protein SPE-18 promotes localized assembly of the major sperm protein in *C. elegans* spermatocytes

**DOI:** 10.1101/2020.08.10.244988

**Authors:** Kari L. Price, Marc Presler, Christopher M. Uyehara, Diane C. Shakes

**Affiliations:** Department of Biology, William & Mary, Williamsburg, VA 23187, USA; Department of Genetics, Yale University School of Medicine, New Haven, CT; Applied BioMath LLC, Concord, MA; Department of Genetics, The University of North Carolina at Chapel Hill, Chapel Hill, NC

**Keywords:** major sperm protein, intrinsically disordered protein, *Caenorhabditis elegans*, spermatogenesis, cytoskeletal assembly, *spe-18*

## Abstract

Many specialized cells use unconventional strategies of cytoskeletal control. Nematode spermatocytes discard their actin and tubulin following meiosis, and instead employ the regulated assembly/disassembly of the Major Sperm Protein (MSP) to drive sperm motility. However prior to the meiotic divisions, MSP is effectively sequestered as it exclusively assembles into paracrystalline structures called fibrous bodies (FBs). The accessory proteins that direct this sequestration process have remained mysterious. This study reveals SPE-18 as an intrinsically disordered protein that that is essential for MSP assembly within FBs. In *spe-18* mutant spermatocytes, MSP remains cytosolic, and the cells arrest in meiosis. In wildtype spermatocytes, SPE-18 localizes to pre-FB complexes and functions with the kinase SPE-6 to recruit MSP. Changing patterns of SPE-18 localization revealed unappreciated complexities in FB maturation. Later, within newly individualized spermatids, SPE −18 is rapidly lost, yet SPE-18 loss alone is insufficient for MSP disassembly. Our findings reveal an alternative strategy for sequestering cytoskeletal elements, not as monomers but in localized, bundled polymers. Additionally, these studies provide an important example of disordered proteins promoting ordered cellular structures.

**Summary Statement:** Intrinsically disordered proteins are increasingly recognized as key regulators of localized cytoskeletal assembly. Expanding that paradigm, SPE-18 localizes MSP assembly within *C. elegans* spermatocytes.

## INTRODUCTION

The ability of cells to move, divide, and assume specific cell shapes requires a cytoskeleton that can reversibly assemble into a wide range of structures. Core to this flexibility is the intrinsic capacity of core molecular subunits to polymerize into filaments. The subsequent process of regulating how, when, and where these filaments assemble into larger molecular superstructures is directed by a wide diversity of modifier and accessory proteins (Hohmann and Dehghani, 2019; Rottner et al., 2017; Goodson and Jonasson, 2018). Current concepts of cytoskeletal regulation have been dominated by functional studies of actin and tubulin and their interactions with diverse accessory proteins (Svitkina, 2018; Buracco et al., 2019; Brouhard and Rice, 2018; Bodakuntla et al., 2019; de Forges et al., 2012). However, a full understanding of cytoskeletal control requires that we also consider less-studied proteins whose properties challenge our standard assumptions.

One such protein is the nematode Major Sperm Protein (MSP), whose assembly/disassembly dynamics power the crawling motility of nematode spermatozoa (Klass and Hirsh, 1981; Sepsenwol et al., 1989; Italiano et al. 1996; and reviewed in Roberts and Stewart, 2012; Smith, 2014). Although MSP-based motility appears superficially similar to its actin-based counterpart, the molecular mechanisms are distinct. Much of what we know about MSP dynamics was gleaned from the parasitic nematode Ascaris, whose size and sperm number make Ascaris sperm amenable for biochemical studies. MSP lacks nucleotide binding sites and is quite small, only 14kDa (Roberts, 2005). Importantly, while polarity is a hallmark of actin and tubulin assembly, MSP monomers form symmetric homodimers that subsequently form apolar filaments (Bullock et al., 1998). Because MSP filaments lack polarity, they are not associated with molecular motors, and their unidirectional growth requires accessory proteins. *In vitro* comet assays show that the integral membrane protein MPOP is sufficient for MSP polymerization (LeClaire et al., 2003). However, within crawling spermatozoa, the localized assembly of MSP filaments involves several additional factors including a serine/threonine (ser/thr) kinase MPAK, a filament assembly factor MFP2 that is activated by MPAK, a growing end capping protein MFP1, and a filament stabilizing factor MFP3 (Roberts and Stewart, 2012). Disassembly of MSP filaments at the base of the pseudopod involve dephosphorylation of MFP3 by a PP2A phosphatase (Yi et al., 2009).

Non-flagellated, crawling spermatozoa are a defining feature of the phylum Nematoda, and these MSP-propelled cells are both remarkably speedy (Italiano et al., 1999) and highly efficient; in the hermaphroditic species *Caenorhabditis elegans*, every sperm successfully fertilizes an oocyte (Singson, 2001). Yet the developmental program required to produce these spermatozoa includes both assets and challenges. In *Caenorhabditis elegans* where it has been best studied, spermatogenesis occurs in a linear developmental sequence along the length of the gonad (Fig. 1A). Instead of taking days to weeks as in Drosophila and vertebrates, progression through the stages of meiotic prophase takes less than 24 hours (Jarmillo-Lambert et al., 1999; Fig. 1A, C, D), and post-meiotic development is abbreviated to minutes rather than days (Chu and Shakes, 2013). Two key factors account for the brevity of the post-meiotic process. First, instead of having to remodel their actin and tubulin into specialized structures following the meiotic divisions, nematode spermatocytes discard their actin and tubulin into a central residual body and MSP takes over as the core cytoskeletal element in haploid sperm (Nelson et al., 1982; Ward, 1986; Winter et al., 2017; Fig. 1E,F). Second, during meiotic prophase, nematode spermatocytes must synthesize and pre-package all of the components needed to support post-meiotic sperm functions. Global transcription ceases near the end of meiotic prophase, precluding any post-meiotic burst of sperm-specific transcription (Shakes et al., 2009); and protein synthesis ceases as the cell’s ribosomes are discarded into the residual body (Ward et al, 1981). These efficiencies are countered by the challenge of how to control the potentially disruptive random self-assembly of MSP as MSP levels rise to compromise 10-15% of the total and 40% of the soluble cellular protein (Roberts, 2005).

**Figure 1.**
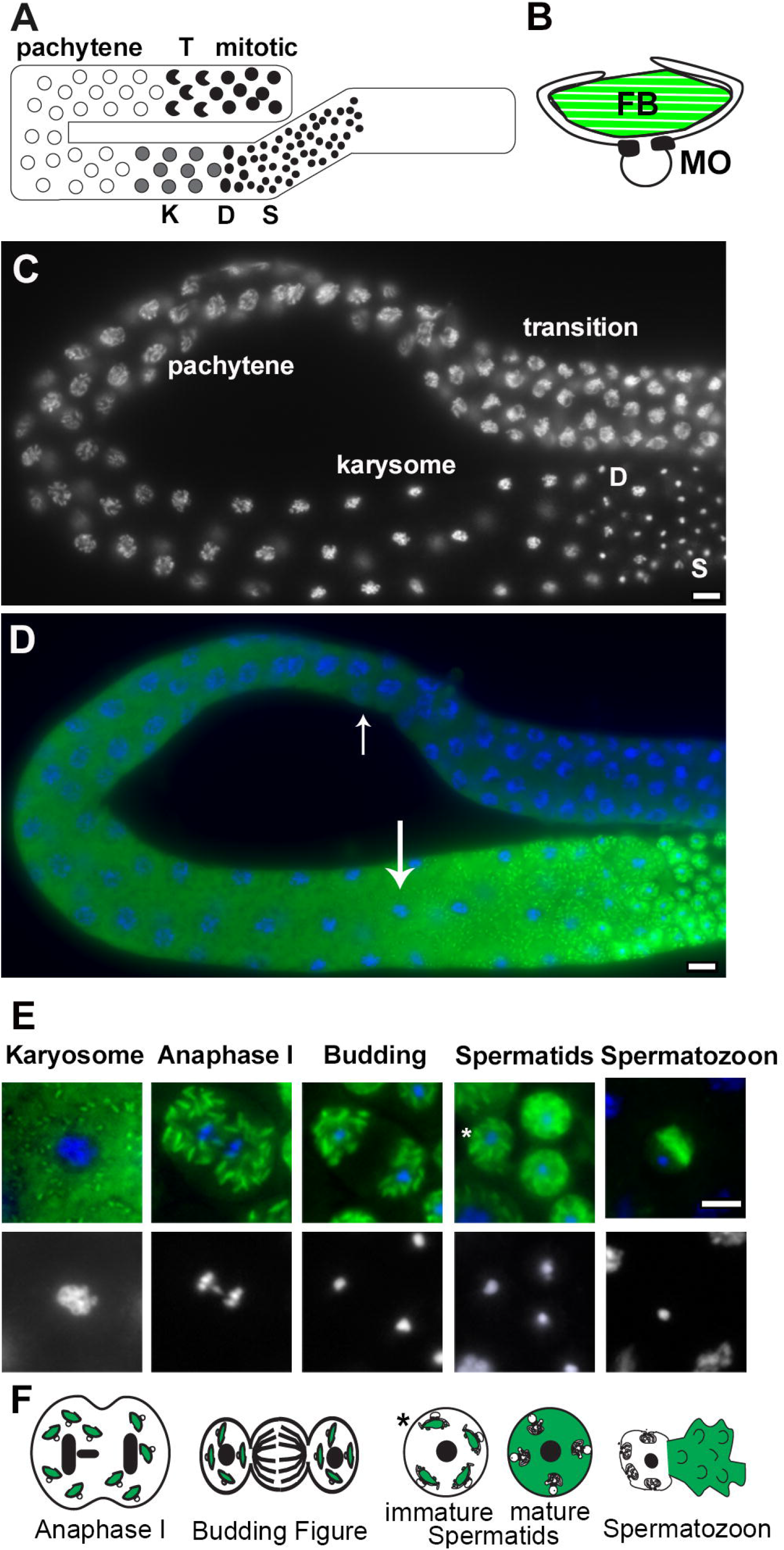
Overview of *C. elegans* spermatogenesis. (A) Schematic of adult male gonad highlighting its linear organization. Undifferentiated germ cells proliferate mitotically at the distal end, then commit to spermatogenesis as they transition (T) to meiotic prophase before entering an extended pachytene stage. Towards the end of meiotic prophase, the spermatocytes enter the karyosome stage (K) during which the chromosomes compact and global transcription ceases. Following a narrow zone of meiotically dividing spermatocytes (D), quiescent haploid spermatids (S) accumulate in the seminal vesicle. (B) Schematic of a Golgi-derived fibrous body-membranous organelle (FB-MO) complex showing the arms of the MO head region surrounding the MSP-enriched FB (green), the glycoprotein filled MO vesicle, and the electron dense collar that divides these two domains of the MO. (C-D) Isolated male gonad showing stage-specific chromatin morphology by DAPI (C) and co-labelled with anti-MSP (green) to show initial expression in pachytene spermatocytes (small arrow) and distinct FBs (large arrow) in karyosome stage spermatocytes (D). (E-F) Stage-specific patterns of MSP distribution in spermatocytes co-labelled with DAPI (blue) and anti-MSP (green) or in schematic drawings (F). During nematode spermatogenesis, anaphase II is following by a partitioning, budding figure stage during which the cell’s actin, microtubules, and ribosomes are discarded in a central residual body while the FB-MO complexes, mitochondria, and chromatin partition to the spermatids. Once spermatids detach from the residual bodies, all but the most recently individualized (*), contain MOs that have docked but do not fuse with the plasma membrane and the FBs disassemble so that MSP disperses throughout the cytoplasm. In motile spermatozoa, the MOs form stable fusion pores with the plasma membrane of the cell body, and MSP localizes to the pseudopod where its assembly/disassembly dynamics drive pseudopod motility. Scale bars = 5 microns.

Developing spermatocytes address this challenge by assembling MSP into a distinct, stable, and sequestered form (Fig. 1). Little is known about the accessory proteins that govern this alternate mode of MSP assembly. However, imaging in *C. elegans* reveals the following sequence. MSP is first detectable in the cytosol of spermatocytes during meiotic prophase, specifically in mid-pachytene spermatocytes when other sperm function proteins are first synthesized (Chu and Shakes, 2013; Fig. 1C,D). Then, towards the end of meiotic prophase (karyosome stage), MSP packs into symmetrically elongating structures called fibrous bodies (FBs) (Fig 1B). These individual FBs develop in close association with Golgi-derived organelles known as a membranous organelles (MOs) (Roberts et al., 1986; Fig. 1B). These FBs are filled with parallel 4.5 nm filaments (Roberts et al, 1986) that contrast with the 11 nm diameter filaments involved in sperm motility (King et al., 1994; Bullock et al., 1998). As MSP is synthesized, its localized polymerization at FBs promotes both FB growth and MSP sequestration. MSP remains locked in these FB structures through the post-meiotic partitioning process during which FB-MO complexes partition to individual spermatids and away from the central residual body (Fig. 1E-F). Once spermatids detach from the residual body, the FB-MO complexes disassociate, the MOs dock with the plasma membrane, and the FBs disassemble into MSP dimers (Roberts et al., 1986).

The packing of MSP into FB-MO complexes is hypothesized not only to prevent MSP from interfering with the actin and tubulin mediated events of meiotic chromosome segregation and cell division (Chu and Shakes, 2013) but also to facilitate MSP partitioning to spermatids during the post-meiotic budding division (Nishimura and L’Hernault, 2010, Fig. 1E-F). However, the necessity of MSP sequestration has never been directly addressed. Additionally, little is known about the composition of FBs. They are assumed to consist solely or largely of MSP, but in principle would require their own set of accessory proteins, like those required to mediate MSP-mediated motility.

Here, we identify *spe-18*, a gene identified in a screen for <u>spe</u>rmatogenesis-defect mutants, as an essential factor in nematode spermatogenesis and FB assembly. In the absence of SPE-18, MSP remains cytosolic rather than assembling into FBs, and no haploid sperm are produced as the developing spermatocytes arrest without undergoing proper meiotic divisions. We show that the *spe-18* gene encodes an intrinsically disordered protein, whose subcellular localization pattern within wild type and mutant spermatocytes suggests that it functions to both localize and structure FB assembly.

## RESULTS

### *spe-18* (*hc133*) mutants produce arrested spermatocytes with cytosolic MSP

Until recently, the only factor known to be required for the initial assembly of MSP into FBs was the ser/thr kinase SPE-6. In *spe-6* mutant spermatocytes, MSP remains cytosolic, and the spermatocytes arrest development without completing the meiotic divisions or undergoing cytokinesis (Varkey et al., 1993; Muhlrad and Ward, 2002; Fig 2A). To identify other factors required for the assembly of MSP into FBs, we examined other spermatocyte arrest mutants for defects in MSP assembly. One proved to be the early acting spermatogenesis-specific transcription factor *spe-44* (Kulkarni et al., 2012), while the other was *spe-18*(*hc133*), previously annotated as *spe-7* (Kulkarni et al., 2012; Chu and Shakes, 2013) and originally isolated in a screen for <u>spe</u>rmatogenesisdefective mutants by D. Shakes and S. L’Hernault (Fig. 2A). To further characterize *spe-18*(*hc133*) mutants, we first confirmed that they exhibited the standard characteristics of SPE mutants; namely that mutant hermaphrodites produce few or no self-progeny but produce cross-progeny when mated to wildtype males (L’Hernault et al., 1988; Nishimura and L’Hernault, 2012). This result indicates that sperm not oocytes are responsible for the fertility defect. To determine if the mutation was temperature-sensitive, we analyzed the self-fertility of mutant hermaphrodites at three temperatures (Table 1). In every case, control hermaphrodites produced >100 progeny and a small number of unfertilized oocytes. These brood sizes are lower than wildtype but reflect the lower fertility related to both the *unc-4* morphological marker and the *him-8* (high incidence of males) mutation that used to increase the number of males. In contrast, *spe-18* hermaphrodites produced no embryos and laid only a small number of unfertilized oocytes. While most temperature-sensitive mutants exhibit more severe defects at elevated temperatures, the self-fertility defect of *spe-18* hermaphrodites was mildly cold-sensitive; *spe-18* hermaphrodites were completely infertile at 16°C, but at 25°C, they laid more unfertilized oocytes and produced as many as eight offspring. In no case did we detect dead embryos, suggesting that when fertilization-competent sperm were produced, they generated viable offspring.

**Figure 2.**
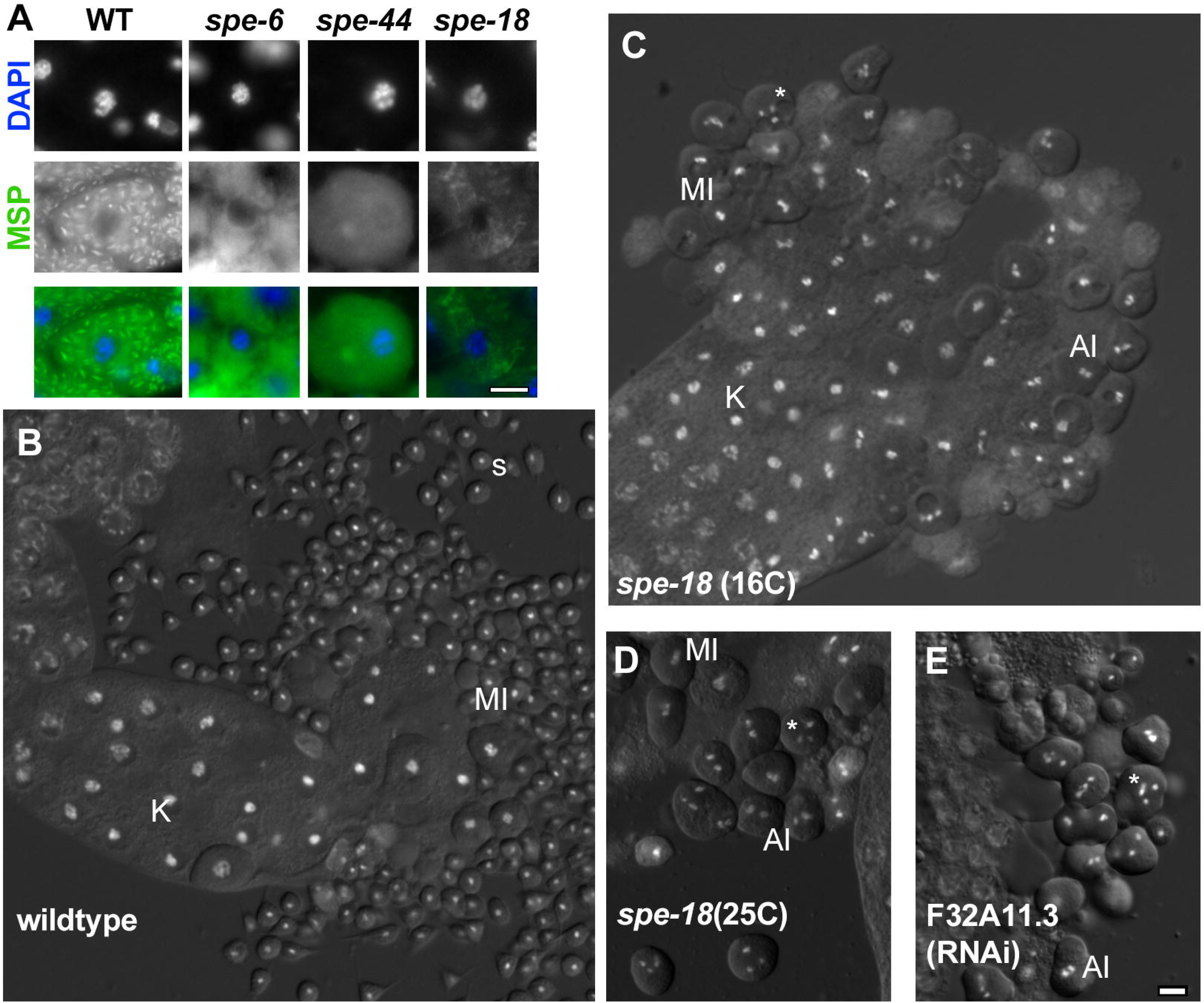
*spe-18* spermatocytes are defective in FB assembly and progression through the meiotic divisions. (A) Wildtype and mutant late meiotic prophase (karyosome) stage spermatocytes co-labelled with DAPI (blue) and anti-MSP (green) and enlarged 1.5X. (B-E) DIC/Hoechst image of wildtype (B), *spe-18* at 16°C (C) and 25°C(D), and F32A11.3 (RNAi) (E) sperm spreads. Abbreviations: karyosome (K), metaphase I(MI); anaphase I(AI), and haploid spermatids (s). Asterisk marks arrested spermatocytes with 3-4 compact chromosome masses. Scale bars = 5 microns.

**Table 1:** Analysis of Hermaphrodite Self-Sterility Phenotype.

| STRAIN | Temp (C) | n | Oocytes (SE) | Embryos (SE) | %Hatch | Progeny (SE) |
| --- | --- | --- | --- | --- | --- | --- |
| *unc-4; him-8* | 16 | 8 | 18.3 +/-5.9 | 183.8+/-2.9 | 87% | 160+/-2.6 |
|  | 20 | 9 | 72.2+/-4.8 | 187.9+/-9.9 | 90% | 170+/-9.8 |
|  | 25 | 7 | 12.9+/-2.6 | 135.1+/-6.0 | 73% | 99.6+/-3.3 |
| *spe-18 unc-4;* | 16 | 22 | 9.6+/-2.0 | 0 | n/a | 0 |
| *him-8* | 20 | 20 | 17.05+/- 1.4 | 0.7+/-0.2 | 100% | 0.7+/-0.2 |
|  | 25 | 17 | 47.6+/-4.3 | 2.4+/-0.5 | 100% | 2.4+/-0.5 |

\*S.E. indicates standard error of the mean.

Analysis of isolated and flattened male gonads revealed that *spe-18* spermatocytes have defects in both meiotic chromosome segregation and cytokinesis. Control gonads included spermatocytes at all stages of development including a small number of meiotically dividing spermatocytes and large numbers of round, haploid spermatids (Fig. 2B). In contrast, *spe-18* gonads lacked haploid spermatids and instead accumulated large numbers of spermatocytes that were the size of primary spermatocytes (Fig. 2C-D). Like the hermaphrodite self-fertility, the relative severity of the meiotic chromosome segregation defects was also mildly cold-sensitive. Although most of these chromosome segregation phenotypes were observed at all temperatures, mutant spermatocytes most typically arrested with a single chromatin cluster at 16°C (Fig. 2C), two chromatin clusters at 20°C (data not shown), and 3-4 chromatin clusters at 25°C (Fig. 2D). With the exception of a few spermatocytes at 25°C, spermatocytes failed to undergo either the standard myosin II based cytokinesis following anaphase I or the distinct myosin VI based budding division that normally follows anaphase II (Ward et al., 1981; Winter et al., 2017; Hu et al., 2019).

### SPE-18 is conserved in diverse nematodes and is predicted to contain extended intrinsically disordered regions

To better understand the molecular role of SPE-18 in spermatogenesis, we first needed to clone the *spe-18* gene. We mapped the *hc133* mutation to a small region of chromosome II; and of 43 genes within this interval, only one gene, F32A11.3, had been previously identified in large-scale microarray studies as exhibiting a “spermatogenesis-enriched” expression pattern (Reinke et al., 2000; Reinke et al., 2004). To determine whether the F32A11.3 gene in *spe-18* mutants contained a molecular lesion, we amplified and sequenced the F32A11.3 gene from wildtype and *spe-18* (*hc133*) worms and found that *hc133* contains a C/T point mutation in the last exon that changes the glutamine (Q301) CAA codon to the premature stop codon TAA (Fig. 3A).

**Figure 3.**
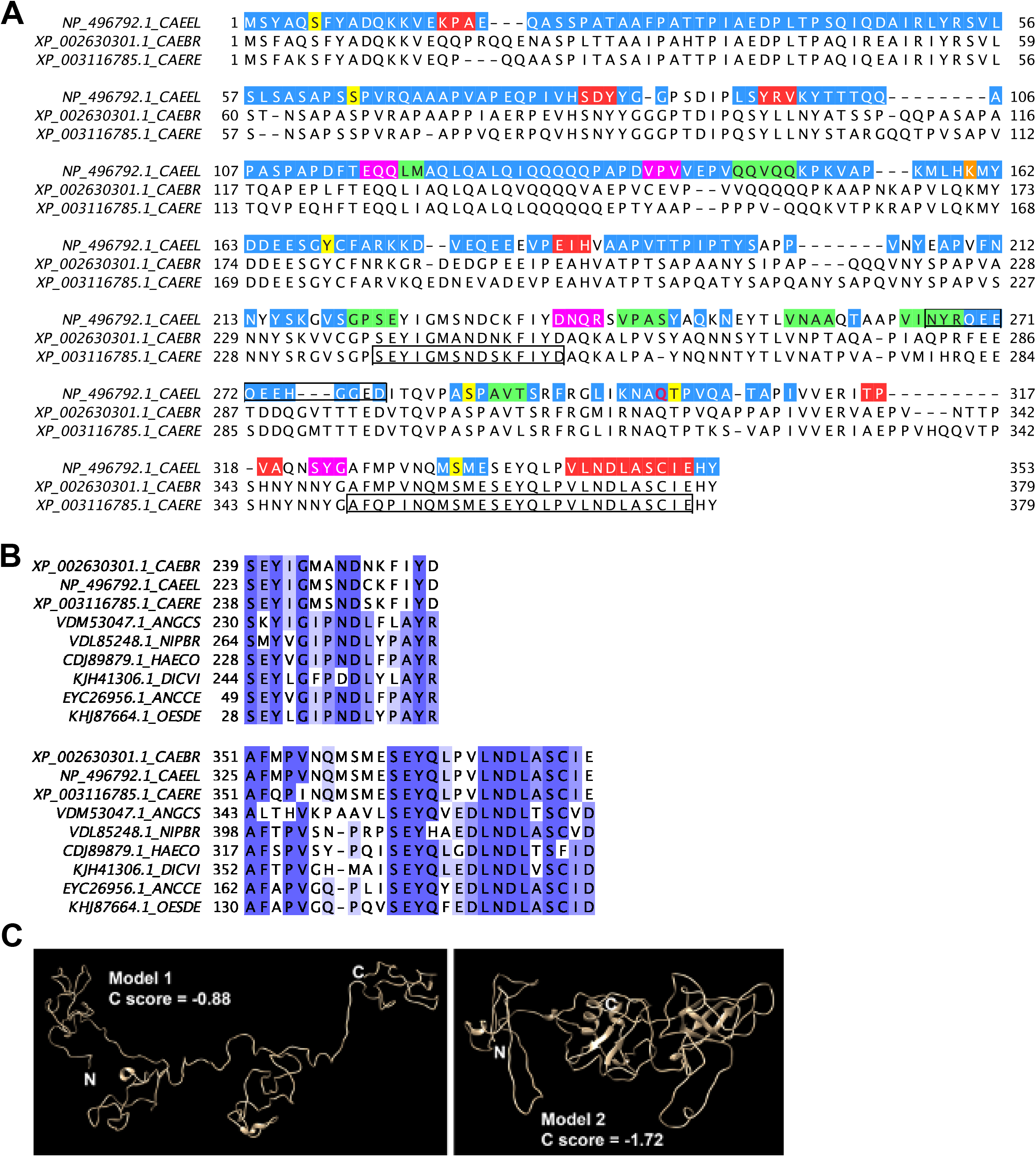
The amino acid sequence of SPE-18 (F32A11.3) and its bioinformatic analysis. (A) Clustal Omega alignment (Madeira et al., 2019) of F32A11.3 with *C. briggsae* and *C. remanei*. Pre-mature stop codon in *hc133* (CAA/TAA) marked in red (Q301). Predicted disordered regions from Phyre2 in blue (Kelley et al., 2015). iTasser structural predictions from model 1 (magenta helices) or model 2 (red helices, green strands) (Roy et al., 2010). Potential phosphorylation sites that are both conserved and predicted with high confidence by NetPhos3.1 highlighted in yellow (Blom et al., 1999). High confidence predicted ubiquitination site from UbPred highlighted in orange (Radivojac et al., 2010). Boxed region (266-279) in the *C. elegans* sequence is the peptide used to generate an antibody. Boxed regions in the *C. remanei* sequence are conserved across multiple nematode species. (B) Regions of high conservation across multiple nematode species corresponding to the boxed regions in A. Species include *Caenorhabditis elegans* (CAEEL), *Caenorhabditis briggsae* (CAEBR), *Caenorhabditis remanei* (CAERE), *Angiostrongylus costaricensis* (ANGCS), *Nippostrongylus brasiliensis* (NIPBR), and *Ancylostoma ceylanicum* (ANCCE) from the order Rhabditida as well as *Haemonchus contortus* (HAECO), *Dictyocaulus viviparus* (DICVI), and *Oesophagostomum dentatum* (OESDE) from the order Strongylida. (C) Top two-scoring iTasser models of SPE-18 protein structure.

To verify that F32A11.3 encoded *spe-18*, we used RNAi feeding to deplete F32A11.3 in *him-8* hermaphrodites and their male progeny. F32A11.3 depleted males exhibited spermatocyte defects that were visually indistinguishable from those of *spe-18*(*hc133*) males (Fig. 2E). Together, these results confirmed the molecular identity of *spe-18*. Furthermore, since RNAi knockdowns invariably represent loss-of-function phenotypes, the RNAi phenotype suggests that the truncation of SPE-18 in *spe-18*(*hc133*) mutants represents a loss-of-function, rather than a neomorphic phenotype.

*spe-18* encodes a 353 amino acid protein (Fig 3A) that lacks any known functional domains. BLASTP analysis identified highly conserved homologs of F32A11.3 within multiple members of the *Caenorhabditis* genus (Fig. 3A). A BLASTP search to nematodes outside of the *Caenorhabditis* genus revealed homologs in species from the larger *Rhabditida* order as well as the order *Strongylida* (Fig. 3B, S1). Alignments to these less conserved homologs revealed two extended regions of high sequence conservation, one central and near the C-terminus, as well as shorter regions of conservation throughout (Fig. 3B, S1).

Multiple lines of evidence from amino acid composition, bioinformatics, and biochemistry suggest that SPE-18 is largely unstructured. The amino acid composition itself reveals that SPE-18 is an acidic protein with an isoelectric point of 4.78. The protein is rich in the disorder-promoting residues proline (P), glutamine (Q), glutamic acid (E), and serine (S), but it also has abundant alanines (A) and valines (V) (Fig. 3A). Bioinformatic studies show that SPE-18 lacks transmembrane domains, and two distinct disorder predicting programs suggest that SPE-18 has large intrinsically disordered regions. Phryre2 (Kelley et al., 2015) predicts that it is 70% unstructured (Fig. 3A). PrDOS (Ishida and Kinoshita, 2007) predicts that SPE-18 contains 25 to 50% unstructured residues depending on the false positive setting; these amino acids were largely a subset of those identified by Phrye2. In addition, the most likely model predicted by structure modeling program iTasser (Roy et al., 2010) suggests that SPE-18 possesses minimal secondary structure (Fig. 3A, C). In this context, it is notable that both iTasser model 2 and PSSpred (Yan et al., 2013) predict that the conserved C-terminal domain contains a ten amino acid alpha helix (Fig. 3A,C). Furthermore, when this C-terminal region is deleted in *hc133* mutants; the truncated protein is destabilized. Finally, one key biochemical property of intrinsically unstructured proteins is that they are heat stable (Uversky, 2017). To test the inherent heat-stability of SPE-18, we expressed recombinant SPE-18 in *E. coli* and then assayed whether SPE-18 within the resulting lysate remained in the supernatant after a ten minute heat treatment at 95°C. Under these conditions, most proteins within the lysate precipitated whereas SPE-18 remained in the supernatant (Fig. S2). Collectively, these data predict that SPE-18 functions as an intrinsically disordered protein.

As the function of intrinsically unstructured proteins is often regulated by post-translational modifications, we also employed to bioinformatic approaches to assess potential phosphorylation sites. NetPhos3.1 predicted several high confidence phosphorylation sites in SPE-18 that are also conserved in its *Caenorhabditis* homologs (Fig. 3A) and two (S6 and Y169) that are conserved in more distant species (Fig. S1).

Taken together, these data predict that SPE-18 functions as a protein with large intrinsically disordered regions. However the sequence alignments also indicate that SPE-18 contains both extended and shorter conserved regions that could potentially serve as sites either for molecular interactions or for regulation by post-translational modifiers.

### SPE-18 Protein Localizes in a Stage-Specific Pattern to FBs of Developing Spermatocytes

To understand how SPE-18, as an unstructured protein, was promoting the assembly of MSP into fibrous bodies (FBs), we next sought to determine the cellular distribution of SPE-18. Does SPE-18 direct localized MSP assembly as a resident protein of either the FB or MOs, or does SPE-18 direct FB assembly from some other cellular compartment? Is SPE-18 only present in spermatocytes or might it also be present in haploid sperm such that it could regulate MSP function at multiple stages of spermatogenesis?

To address these questions, we first generated polyclonal antisera to a region of SPE-18 that was predicted to be both antigenic and specific (Fig. 3A). Since the antigenic sequence is before the *hc133* truncation, the antibody was predicted to recognize both the full-length and truncated protein. Western blots were used to test the specificity of the anti-SPE-18 antibody (Fig. 4A). Anti-SPE-18 antibody bound to a 42 kDa protein in lysates of wildtype adult males but not in *spe-18* (*hc133*) males or males lacking an essential transcription factor for *spe-18, spe-44*(*ok1400*) (Kulkarni et al, 2012; Fig. 4A). This result not only confirmed the specificity of the antibody but also revealed that the *hc133* allele is functionally null as no truncated protein could be detected.

**Figure 4.**
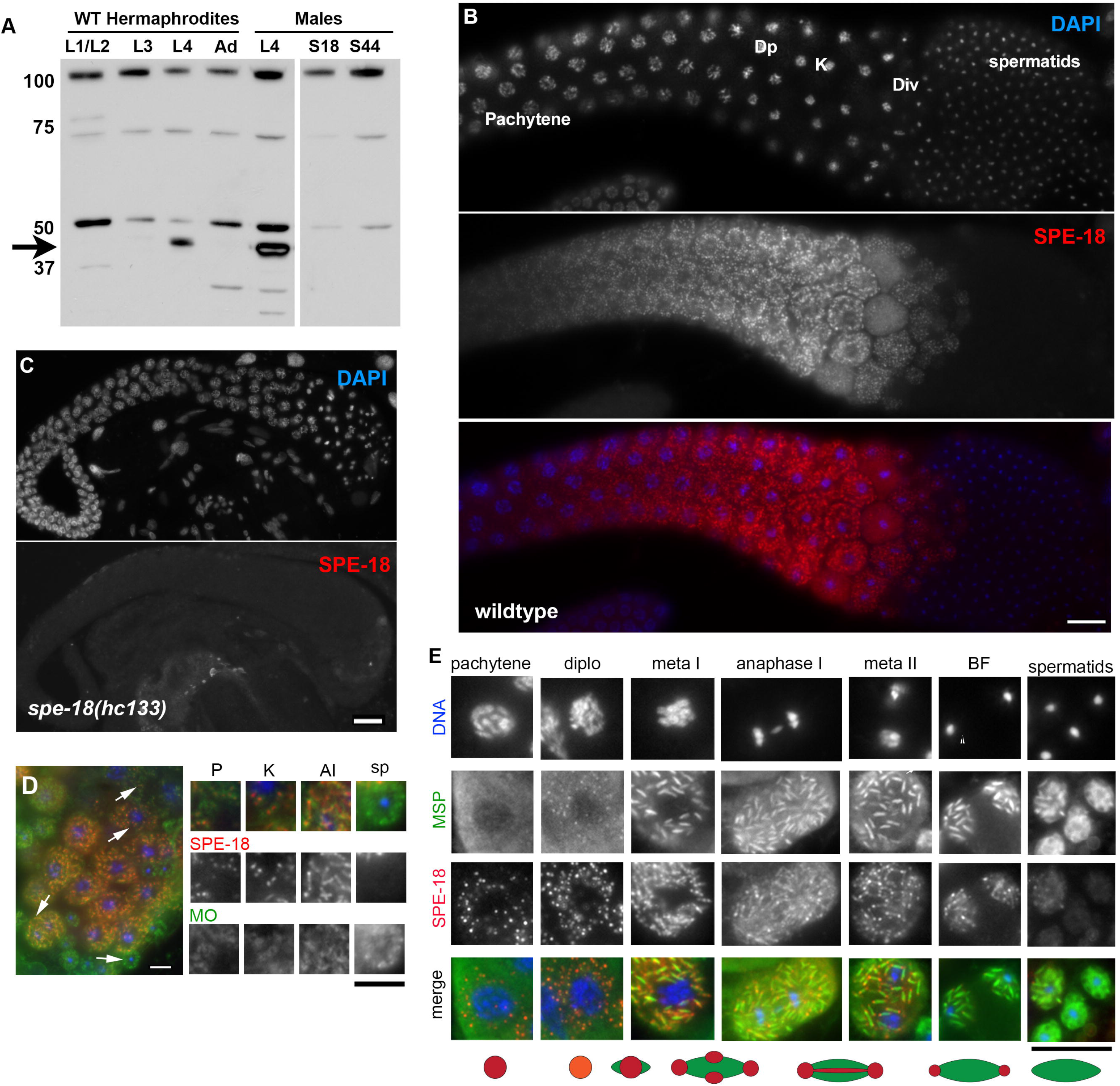
SPE-18 localizes to developing FBs in a stage-specific pattern. (A) Western blot comparing SPE-18 levels in age-synchronized wildtype hermaphrodites, wildtype males, and mutant *spe-18* (S18) and *spe-44* (S44) males. A non-specific band at ~100 kDa serves as a loading control. Arrow in shows the position of SPE-18 with strong bands in L4 hermaphrodites and males. (B,C) Immunocytology of isolated male gonads co-labelled with DAPI and anti-SPE-18 antibody in wildtype males (B) or *spe-18*(*hc133*) males (C). (D) Co-immunolabelling of wildtype spermatocytes with anti-SPE-18 and the MO marker 1CB4. Inserts are 3X enlarged for visibility and correspond to arrows in the larger figure. (E) SPE-18/MSP co-immunolabelling in individual, staged, wildtype spermatocytes and cartoon schematic. 2X enlarged. Abbreviations: Pachytene (P), Diplotene (Dp/Diplo), karyosome (K), meiotic division zone (Div), metaphase I/II (meta I/II); budding figure (BF), anaphase I (AI), spermatids (sp). Scale bars = 5 microns (D) and 10 microns (B, C, E).

On the same western blot, we tested hermaphrodite samples from specific larval stages (Fig 4A) and found that the major band detected in adult males could only be detected in fourth stage larvae (L4), the only stage when hermaphrodites are actively producing sperm. The notable absence of SPE-18 in adult hermaphrodites that have spermatozoa in their spermathecas, suggested that SPE-18 might function in developing and/or meiotically dividing spermatocytes but not in haploid sperm.

We next determined the subcellular localization of SPE-18 by co-labelling isolated wildtype and mutant male gonads with DAPI and anti-SPE-18 antibody. Within wildtype male gonads, SPE-18 labelling was first detectable in late pachytene spermatocytes and then increased in intensity through the end of the karyosome stage (Fig. 4B). Consistent with the western blots, no signal was detectable within either *spe-18*(*hc133*) (Fig. 4C, Fig. S3C,D) or *spe-44* (Fig. S3E,F) gonads, confirming the specificity of the antibody for immunocytology. Within developing spermatocytes, SPE-18 labelled numerous discrete structures whose pattern and distribution seemed similar to fibrous bodies (FBs) (Fig. 4B). SPE-18 labelling then decreased in intensity through the meiotic divisions and became undetectable in haploid spermatids. This failure to detect SPE-18 in haploid sperm was consistent with the absence of a SPE-18 signal in western blots of adult spermatozoa-containing hermaphrodites. Importantly, since the one key defect in *spe-18*(*hc133*) spermatocytes is the inability to assemble MSP into FBs, clear evidence of SPE-18 localizing to FBs might suggest a direct role for SPE-18 in FB assembly.

Because FBs develop in close association with the Golgi-derived MOs (Fig. 1B), it can be challenging to distinguish between the two compartments. To confirm that SPE-18 is not an MO component, we compared the localization patterns of SPE-18 to the MO marker 1CB4 (Okamoto and Thompson, 1985), a monoclonal antibody which labels multiple MO glycoproteins (Fig. 4D). Within developing pachytene and karyosome stage spermatocytes (Fig. 1), the first detectable SPE-18 structures were adjacent or within the 1CB4 labelled membranes, consistent with the known ultrastructure of FB-MO complexes (Roberts et al., 1986). By anaphase, the SPE-18 labelled FBs were larger than the confines of the MO (Fig. 1B and 4D). In haploid spermatids with MOs docked at the plasma membrane, SPE-18 was undetectable. These results show that SPE-18 is present in spermatocytes but not haploid sperm, and that within developing spermatocytes it associates with MOs during the earliest stages of FB development. Furthermore, the manner in which the MO and SPE-18 patterns diverge as spermatocytes mature suggests that SPE-18 is an early component of the FB rather than of the MO.

If SPE-18 contributes to the localization and/or nucleation of FB assembly, then SPE-18 should localize to developing FBs before MSP. To test this prediction, we compared the localization patterns of SPE-18 and MSP (Fig. 4E) and discovered previously undescribed details of FB growth and morphogenesis. In late pachytene spermatocytes when SPE-18 became detectable in distinct, spherical “pre-FB” structures, MSP was already present but diffuse throughout the cytoplasm. By diplotene when spermatocytes are transitioning from pachytene to the karysome stage, MSP co-localized in spherical structures with SPE-18. Through the karyosome stage, the SPE-18 and MSP patterns diverged such that SPE-18 became differentially enriched at multiple points (typically four) around the edges of each FB. One the spermatocytes were meiotically dividing, their spindle-shaped FBs grew primarily through elongation, with most of the growth seemingly restricted to the two ends. In these elongating FBs, SPE-18 localized in a barbell-like pattern with weak but persistent labelling of a central stripe and high concentrations of SPE-18 at the two ends. FBs reached their maximal size by metaphase II. During the budding division that follows anaphase II, SPE-18 segregated to the spermatids and away from the central residual body. SPE-18 labelling was undetectable in all but the most recently individualized spermatids. In contrast, MSP remained uniformly distributed throughout the FBs until later in the spermatid maturation process when the FBs disassembled, and MSP dispersed throughout the cytoplasm. The final disappearance of SPE-18 correlated with FB disassembly. Conversely the early localization of SPE-18 to pre-FB structures and its subsequent enrichment in regions of FB growth are consistent with SPE-18 functioning to nucleate MSP polymerization and/or promote the growth and bundling of MSP filaments.

During the process of spermatogenesis, cellular components that are no longer needed are typically discarded into the residual body during the post-meiotic budding division (Fig. 1F). Thus we were surprised by the distinct and unusual pattern of SPE-18 partitioning to the spermatids and then becoming undetectable shortly thereafter (Fig. 4B,E). To rule out the possibility that this unusual pattern of SPE-18 loss was an artifact of antigen accessibility, we assessed the relative levels of MSP and SPE-18 by immunocytology and western blots in aging celibate males (Fig. S4). The western blot of sibling males supported our immunocytology results; as males accumulated spermatids, their MSP levels increased while their SPE-18 levels decreased in proportion to the shrinking numbers of late stage spermatocytes. This result confirmed that SPE-18 is indeed lost in newly individualized spermatids.

### In the absence of the kinase SPE-6, SPE-18 still forms nascent pre-FB structures

Since both SPE-18 and the kinase SPE-6 are required for MSP to assemble into FBs, we investigated whether and how SPE-18 localization patterns might be altered in the null mutant *spe-6*(*hc49*) (Muhlrad and Ward, 2002). In *spe-6* spermatocytes, MSP remained uniformly distributed throughout the cytoplasm while SPE-18 localized to discrete “pre-FB” structures (Fig. 5). Similar SPE-18 positive / MSP negative structures are present early, in the pachytene-stage spermatocytes of wildtype males (Fig. 4C). However in *spe-6* spermatocytes, we could only detect these structures in later karyosome stage spermatocytes (Fig. 5B). As *spe-6* spermatocytes progressed toward their terminal pro-metaphase arrest state (Varkey et al., 1993) these SPE-18 structures grew in size but remained as single spherical masses; they did not appreciatively extend or restructure into the multi-point or end-dominated structures observed in wildtype spermatocytes (Fig. 5C,D). These results suggest that the ability of SPE-18 to assemble into pre-FBs structures occurs independently of SPE-6. However SPE-6 is subsequently required either directly or indirectly for MSP and possibly other FB components to add to these pre-FBs. In the absence of normal FB assembly and elongation, SPE-18 fails to reorganize from its initial spherical structures.

**Figure 5.**
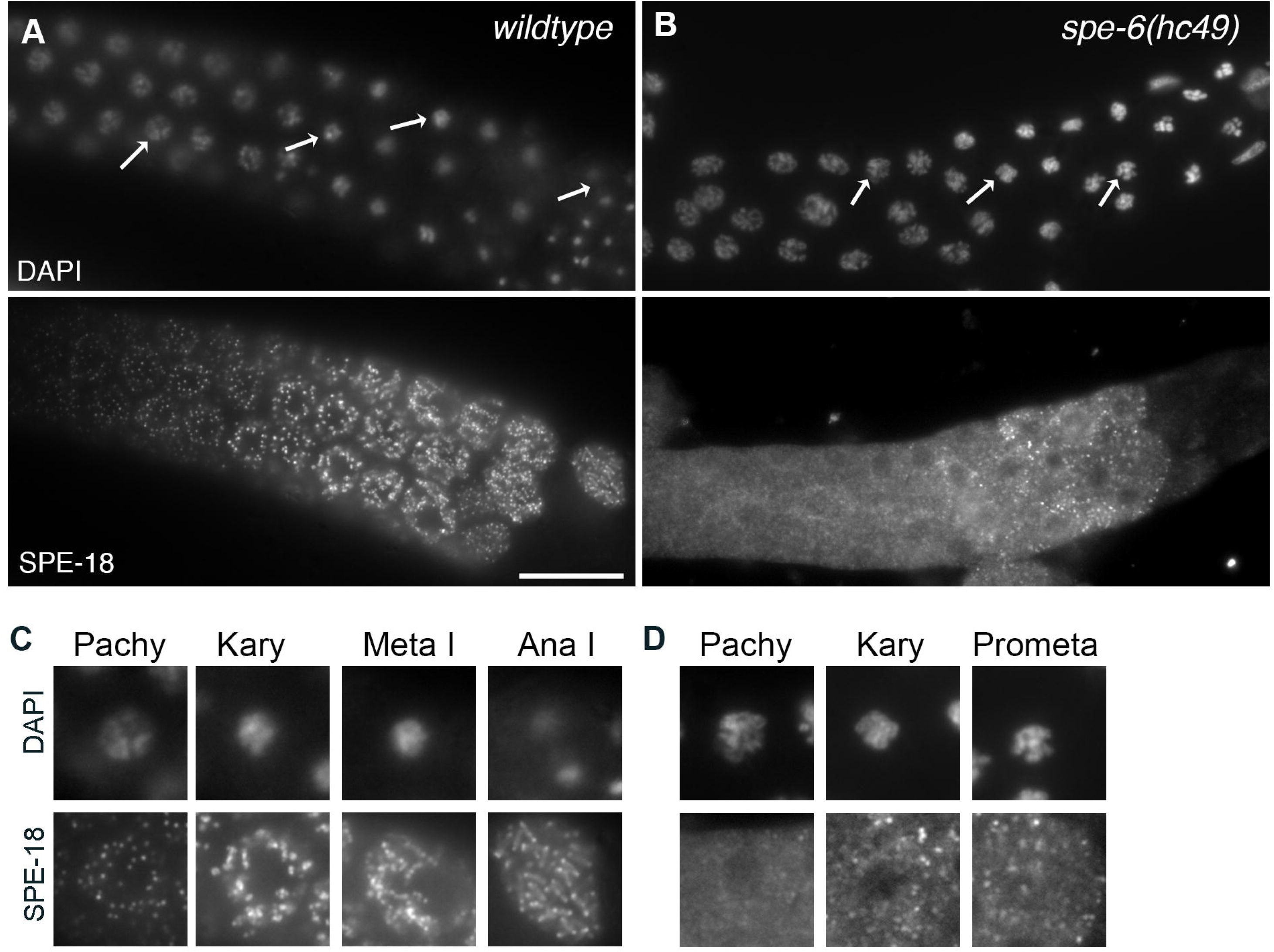
In *spe-6*(*hc49*) mutants, SPE-18 forms pre-FBs in the absence of polymerizing MSP. Immunolocalization of SPE-18 in wildtype (A) and *spe-6*(*hc49*) male gonads (B) co-labelled with DAPI and anti-SPE-18. (C,D) 2X enlarged images of arrow-indicated spermatocytes from gonad image above. Stages include pachytene (pachy), karyosome (kary) and either anaphase I (wildtype) or terminal prometaphase arrest (*hc49*). Scale bar = 20 microns.

### SPE-18 loss is not sufficient for the disassembly of mature FBs

The discovery that SPE-18 concentrates on the ends of mature FBs and that the subsequent loss of SPE-18 correlates with FB disassembly suggested that SPE-18 might not only promote nascent FB assembly within developing spermatocytes but SPE-18 might also serve a later capping and/or stabilizing function in mature FBs. However if SPE-18 does play an essential role in stabilizing the ends of mature FBs then we would expect SPE-18 to persist in spermatids in which FB disassembly is blocked or delayed. To test this hypothesis, we investigated SPE-18 in two contexts. First, we tested if SPE-18 persisted in mutant spermatids that lacked the sperm-specific P1 phosphatases GSP-3 and GSP-4, as these mutant spermatids maintain much of their MSP in FBs (Wu et al., 2012). However examination of *gsp-3/4* spermatids revealed that SPE-18 loss occurred on schedule, shortly after spermatids detached from residual bodies (Fig. 6A). Next, we examined the spermatids of restrictively grown *fem-3*(*gf*) hermaphrodites which have a female soma but produce only sperm (Barton et al., 1987) since FB disassembly is known to be delayed in these spermatids (Wu et al., 2012). However, in *fem-3*(*gf*) spermatids, we also found that the timing of SPE-18 loss was unaltered (Fig. 6B). Although our results do not address whether the loss of SPE-18 is necessary for FB disassembly, they do indicate that loss of SPE-18 from the ends is not sufficient for disassembly. Instead our results remain consistent with models in which the phosphatases GSP-3/4 promote MSP disassembly (Wu et al, 2012).

**Figure 6.**
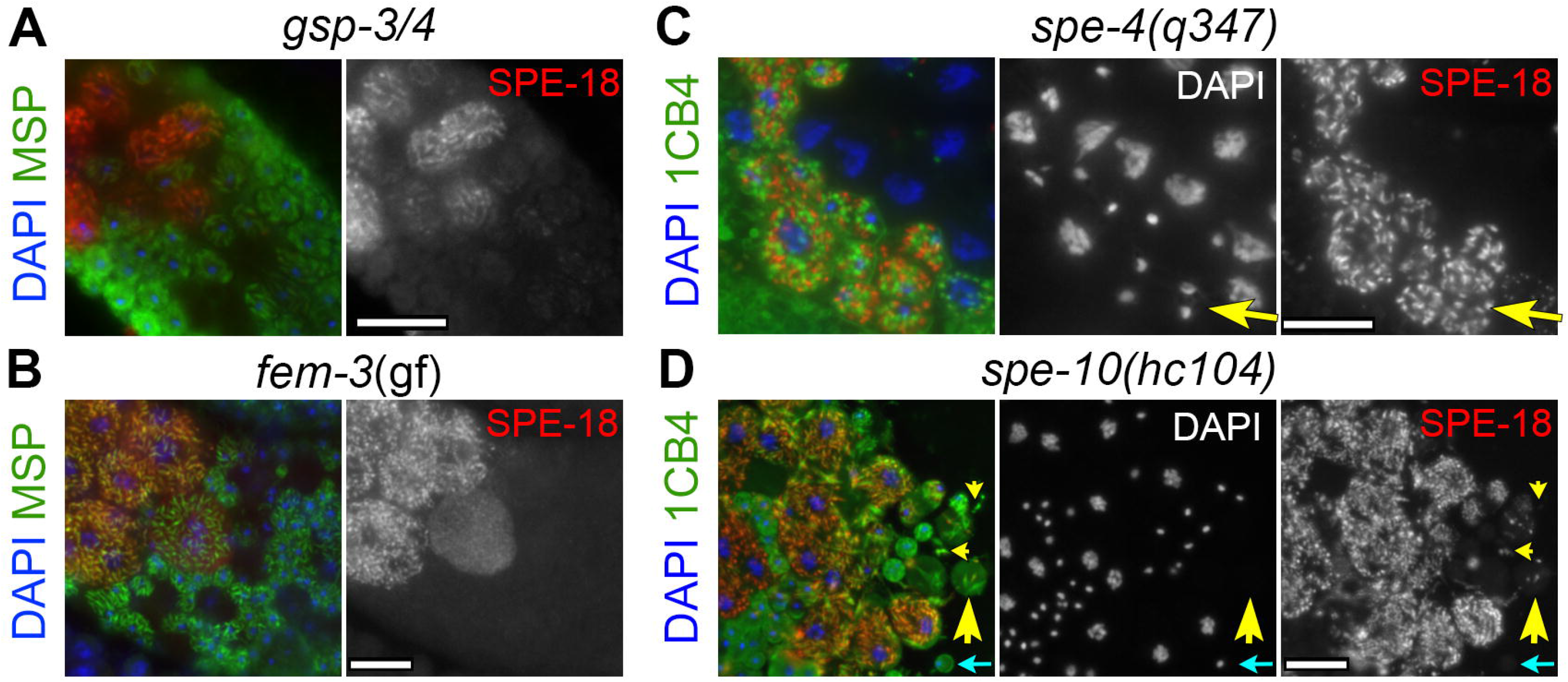
SPE-18 is lost in newly individualized spermatids but stabilized in other cellular contexts. Zone of meiotically dividing spermatocytes, budding figures and spermatids in isolated gonads co-labelled with DAPI (blue), anti-SPE-18 (red), and either anti-MSP (green) (A,B) or the MO marker 1CB4 (green) (D,E). SPE-18 labelling is undetectable in the spermatids of *gsp-3/4* males (A) and *fem-3*(*gf*) (B) hermaphrodites despite the persistent MSP structures. (C) SPE-18 labelling of arrested *spe-4* budding figures (C, yellow arrow). (D) MO marker (1CB4) and SPE-18 labelling of *spe-10* sperm spreads showing FBs in mis-segregated to a residual body (large yellow arrowhead), released FBs (small yellow arrowheads), and spermatids (cyan arrow). Scale bar = 10 microns.

### SPE-18 is stabilized when spermatocytes fail to divide or FBs mis-segregate to the residual body

The rapid disappearance of SPE-18 following sperm individualization raised the question of what regulates the stability of SPE-18. The loss of SPE-18 coincides with several different cellular transitions that could plausibly regulate its degradation. Key among these are 1) the completion meiotic chromosome segregation, 2) the physical separation of the FB from its associated MO, or 3) a physiological difference between spermatocytes and haploid spermatids. In an attempt to rule out some of these possibilities, we first examined SPE-18 patterns in *spe-4* mutants. *spe-4* encodes a presenilin-related, MO transmembrane protein, and mutant spermatocytes complete the meiotic chromosome segregation but fail to complete the budding division (L’Hernault and Arduengo, 1992; Arduengo et al., 1998). In terminally arrested *spe-4* spermatocytes, SPE-18 persisted at elevated levels (Fig. 6C), ruling out a potential linkage to the completion of meiotic chromosome segregation. We next examined mutants in *spe-10*, a palmitoyl transferase and is required for proper partitioning of FB-MOs into spermatids (Shakes and Ward, 1989: Gleason et al., 2006). In *spe-10*(*hc104*) spermatocytes, FBs separate from their MOs prior to spermatid-residual body separation, and a subset of MO-separated FBs either mis-segregate to the residual bodies or form cytoplasts as they bud from the residual bodies (Fig. 6D). Analysis of *spe-10* residual bodies containing FBs revealed that most of these FBs (40/50) labelled with both MSP and SPE-18 antibodies (large yellow arrowhead). For FBs that had budded from the mutant residual bodies as independent cytoplasts, only some labelled with SPE-10 (small yellow arrowheads). SPE-18 was undetectable in *spe-10* spermatids (cyan arrow). Thus, analysis of *spe-10* mutants ruled out a potential linkage to the separation of FB from MOs. Instead this analysis of *spe-4* and *spe-10* spermatocytes favors models in which the loss of SPE-18 is coupled to some property of the individualized spermatids that is distinct from either undivided spermatocytes or residual bodies.

## DISCUSSION

For cells to function properly, polymerization of their cytoskeletal elements must be precisely controlled in both time and space. For many cells, localized polymerization is essential to initiate new cell functions. For nematode spermatocytes, localized MSP polymerization was hypothesized to both package MSP for post-meiotic partitioning and sequester it from interfering the meiotic divisions. In the present study, we show that the spermatogenesis specific protein SPE-18 promotes the localized assembly of the nematode major sperm protein MSP into tightly packed structures known as fibrous bodies (FBs). *spe-18* mutants exhibit sperm-specific sterility, and their spermatocytes are unable to assemble MSP into FBs. Consistent with SPE-18 functioning as a nucleating/assembly factor, SPE-18 is present in the right place at the right time. In wildtype spermatocytes, SPE-18 forms a single spherical “pre-FB” in association with each Golgi-derived membranous organelle (MOs), and these “pre-FBs” form before MSP co-localizes to these structures. SPE-18’s localization is independent of the kinase SPE-6, but SPE-6 is required for MSP to join the pre-FBs. *spe-18* mutant spermatocytes exhibit additional defects in both meiotic chromosome segregation and cytokinesis that are partially cold-sensitive. However, the SPE-18 localization patterns suggest that these defects are likely a secondary consequence of failing to sequester MSP within spermatocytes. We presume that unassembled FB components either physically interfere with these processes and/or induce the spindle-assembly checkpoint.

This analysis of SPE-18 localization patterns suggests a new model of FB assembly (Fig. 7): 1) SPE-18 assembles at each MO to form a spherical pre-FB that functions as a general gathering site for MSP filaments, 2) In a process that requires the kinase SPE-6, MSP is secondarily recruited to these pre-FBs. 3) As MSP levels rise and MSP concentrates at these sites, MSP polymerizes, and the resulting polymers bundle into FBs, 4) As FBs continue to develop, SPE-18 shifts to a multi-point pattern that promotes ongoing FB elongation at the two ends and expansion in the middle. This distribution correlates with the formation of spindle-shaped FBs. 5) As SPE-18 increasingly concentrates at the two ends, the FBs elongate without expanding substantially in width. In this model, SPE-18 both nucleates localized MSP assembly and subsequently shapes the growing FBs by localizing the regions of expansion.

**Figure 7.**
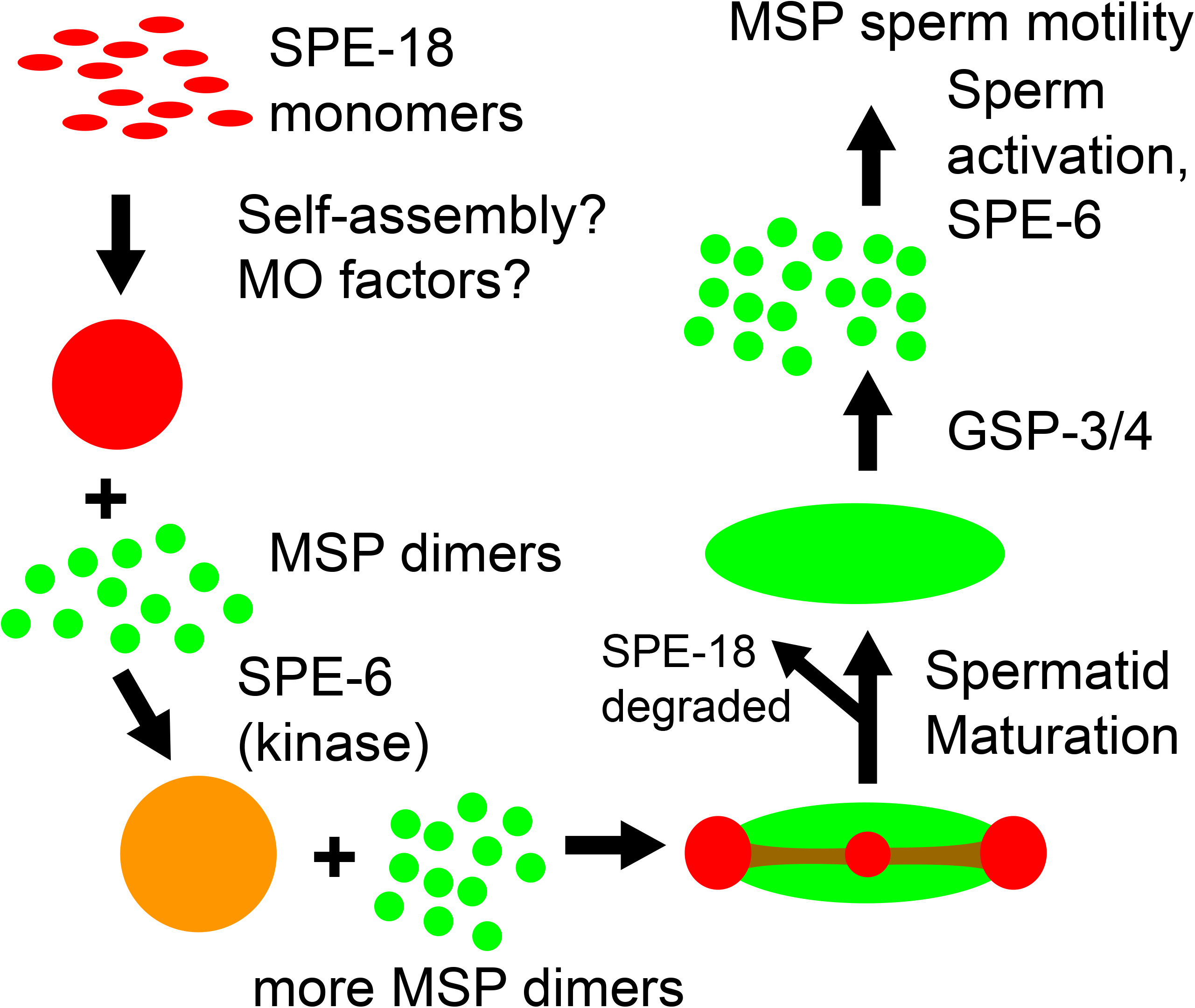
Model of SPE-18 function in localized FB assembly.

Prior to this study, the only known component of FBs was MSP itself. The discovery that the SPE-18 localization patterns change as the FBs develop reveals previously unappreciated complexities in FB composition, growth and shaping. Thus, this study raises new questions regarding both FB composition and control of MSP polymerization. We predict that SPE-18 interacts with multiple binding partners, including diverse FB components and factors that recruit and/or anchor SPE-18 to the MOs While MSP polymerizes differently during FB growth and pseudopod treadmilling, it remains unclear whether this involves distinct or overlapping co-regulators. Although SPE-18 itself is specific to FBs, other components could conceivably function in both contexts. Some of the proteins known to regulate MSP polymerization in the pseudopods of Ascaris sperm (Roberts and Stewart, 2012), may also mediate FB assembly. Candidate interactors are also likely to exist amongst the genes regulated by SPE-44, the transcriptional factor that regulates *spe-18* expression (Kulkarni et al., 2012) or NHR-23, a transcription factor that regulates additional genes required for FB assembly (Ragle et al., 2020).

SPE-18 functions in spermatocytes but is subsequently lost. Rather than being discarded in the residual body, SPE-18 is degraded shortly after differentially partitioning to the sperm. How SPE-18 is lost remains unclear. We found that SPE-18 can be stabilized if it mis-segregates to residual bodies or is trapped within arrested spermatocytes (Fig. 6), so perhaps a SPE-18 stabilizing factor is differentially partitioned to the residual body and away from the sperm. SPE-18, like other intrinsically disordered proteins, could become proteolytically sensitive when released from its binding partners (Uversky, 2017; Flock et al, 2014). Post-translational or pH changes could trigger SPE-18 to disassociate from its binding partners, assume a fully unstructured state, and be subjected to proteolytic degradation. In fact, the SPE-18 sequence includes several potential phosphorylation sites (Fig. 3), and *Ascaris* spermatocytes maintain a higher pH (6.8) than spermatids (6.2) (King et al. 1992; King et al., 1994). SPE-18 does contain a single predicted, high-confidence ubiquitination site K160 (Fig. 3A). However, proteasomes can also degrade intrinsically disordered proteins in the absence of poly-ubiquitination (Asher et al., 2006). A distinct question is why SPE-18 is rapidly degraded following sperm individualization. Perhaps SPE-18 degradation is essential both for FB disassembly and subsequent sperm function. Importantly, SPE-18 loss is insufficient for FB disassembly; as FB disassembly requires the phosphatase GSP-3/4 (Fig. 7; Wu et al, 2012).

The SPE-18 sequence provides important clues regarding how SPE-18 could be promoting FB assembly. SPE-18 is predicted to contain extended intrinsically disordered regions, particularly in the first half of the protein (Fig. 3). SPE-18 also contains two extended highly conserved regions that are not predicted to be disordered (Fig. 3) along with multiple, smaller conserved regions (Fig. S1). These could serve as either binding motifs or sites for post-translational modifications. While the intrinsically disordered regions of SPE-18 are undoubtably critical for its function, they are not sufficient. In the absence of the mostly structured C-terminus, the truncated *hc133* version of SPE-18 is both non-functional and unstable. Proteins with a mix of extended disordered and small structured regions often scaffold the assembly of molecular complexes. Their inherent flexibility paired with multiple high specificity, low affinity binding sites enables them to bind to multiple proteins and exist in multiple distinct conformations (Pancsa and Fuxreiter, 2012). In addition, their ability to rapidly transition between extended and compact conformations enable some to employ a “fly-casting” mechanism to concentrate binding partners. Examples of such proteins include both the actin-modulator Wiskott-Aldrich syndrome protein (WASP) that links cell signaling to localized actin assembly and the phosphatase Calcineurin whose structure facilitates its multi-faceted regulation (Kim et al., 2000; Creamer, 2013). In some cases, the disordered regions of the protein become ordered upon binding to structured proteins (Dyson and Wright, 2002). In others, the disordered regions remain “fuzzy” and never full fold (Sharma et al., 2015).

While the flexibility of proteins with large intrinsically disordered regions can function as singlets to interact with multiple, non-self, binding partners, this class of proteins can also gather together in large assemblages through liquid phase condensation (Shin and Brangwynne, 2017). Within such assemblages, intrinsically disordered proteins may themselves transition from a liquid to solid/amyloid state as they concentrate over time. In other cases, the intrinsically disordered proteins form a liquid droplet within which other proteins, including highly structured proteins, concentrate and polymerize. Examples of this supportive role include the spatial coordination of microtubule nucleation by BuGZ (Jiang et al., 2015), Tau (Hernandez-Vega et al., 2017), PLK4 (Montenegro et al., 2018) and TPX2 (King and Petry, 2020). Notably, the patterns of these protein condensates in association with polymerizing microtubules resembles the patterns we observed of SPE-18 interacting with MSP fibers (Fig. 4E; Fig. 7). In a further parallel, when actin filaments bundle in association with the long flexible cross-linker filamin, the form of the resulting actin superstructures (spheres, spindles, or elongated rods) can be predictably modulated by the filamin-actin ratios (Weirich et al., 2017). Convincing evidence that SPE-18 promotes localized MSP assembly through the process of phase separation awaits both *in vitro* studies and an expanded parts list of FB components. However these intriguing similarities raise the exciting possibility that MSP will join actin and tubulin in the list of cytoskeletal proteins that employ liquid phase condensation to support their localized assembly.

Together these studies have given us new insights into the process and regulation of FB assembly. They place SPE-18 in the context of other known MSP regulators and reveal SPE-18 as an assembly factor for the localized formation and shaping of FBs. Just as studies of MSP assembly/disassembly within the pseudopods of crawling spermatozoa have both challenged and deepened our understanding of actin-based cell motility (Roberts and Stewart, 2000), studies of FB assembly/disassembly dynamics promise to provide an equally informative parallel to our understanding of bundled cytoskeletal structures and their localized assembly. In particular, a deeper understanding of FB dynamics is likely to reveal novel insights into the construction of cytoskeletal assemblages that are facilitated by proteins with large intrinsically disordered regions.

## MATERIALS AND METHODS

### Strains and Culture

*C. elegans* were cultured on MYOB plates (Church et al., 1995) inoculated with *E. coli* strain OP50, using methods similar to those described by Brenner (1974).

Unless otherwise indicated, the following strains were provided by the CGC, which is funded by NIH Office of Research Infrastructure Programs (P40 OD010440):

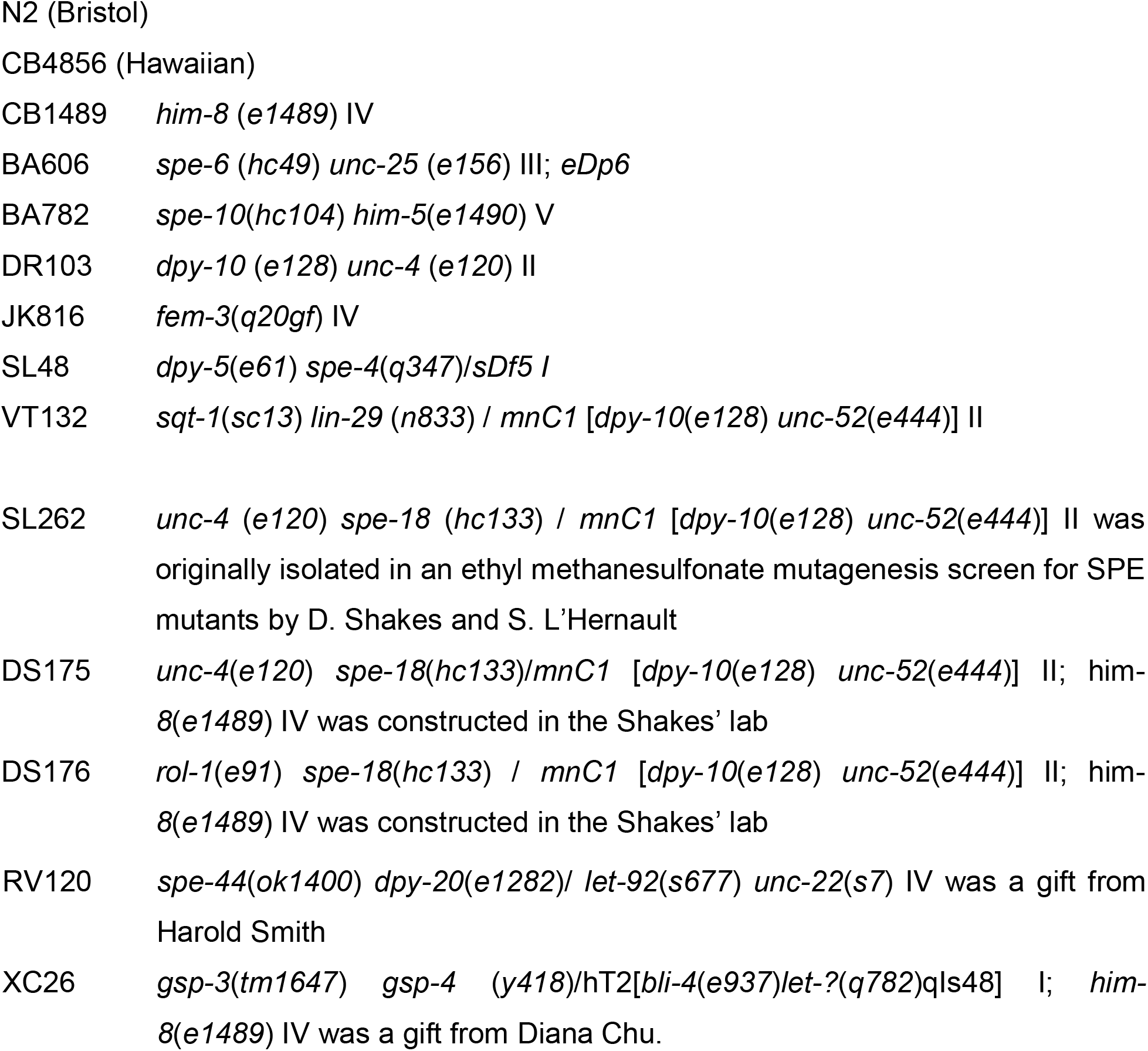

### Fertility analysis

The number of self-progeny for *unc-4; him-8* (wildtype controls) and *spe-18 unc-4; him-8* mutants was determined by placing single worms on separate culture plates and transferring them to fresh plates daily to assess the entire brood.

### Molecular biology, identification, and analysis of the *spe-18* gene

The position of the *spe-18* gene was determined using standard linkage mapping (Sulston and Hodgkin, 1988) and single nucleotide polymorphism (SNP) – mapping (Swan et al, 2002) (see Table S1 in supplementary material). *hc133* was mapped to linkage group II and to the right of *rol-1*. Single nucleotide polymorphisms that generated restriction fragment length polymorphisms (SNIP-SNPs between N2 and Hawaiian (H) strains were used to further position on the physical map. N2/H hybrids were generated by crossing *rol-1*(*e91*) *spe-18*(*hc133*) homozygous hermaphrodites to wildtype Hawaiian males. Rol Non-Spe recombinant offspring from the hybrid worms were isolated and lines were established. Lysates from 18 individual lines and SNP analysis was carried out by PCR amplification using specific primers in the region of the SNP followed by restriction digestion using specific enzymes. Data from this analysis positioned *spe-18* to the right of pkp2112 and close to pkp2116 at approximately 13330 kb. Of the spermatogenesis-enriched genes on linkage group II, F32A11.3 mapped closest to this region.

To identify the molecular lesion in the *spe-18*, the F32A11.3 sequence on Wormbase was used to design primer sets to amplify 500 bp overlapping bidirectional sections. PCR based sequencing was used to sequence F3211.3 from *hc133* mutant DNA in both directions.

For the RNAi experiments, culture plates were soaked with IPTG solution overnight before adding concentrated *E. coli* containing the F32A11.3 feeding construct. Wildtype L4 hermaphrodites were plated on RNAi plates and allowed to lay embryos for 24 hours. The F1 progeny were maintained on these plates and then L4 males were transferred to fresh RNAi plates for an additional 24-48 hours before analysis.

### Immunocytochemistry

To generate anti-SPE-18 antibodies, rabbits were initially pre-screened to identify those whose sera lacked cross-reactivity with *C. elegans* male germlines. Selected rabbits were injected with synthesized peptide corresponding to amino acids 266-279 (YenZym). After a booster injection, serum was collected, and antibodies were affinity purified.

Intact gonads were obtained by dissection of individual males in 5-10 microliters of sperm media (50 mM HEPES, 25 mM KCl, 1 mM MgSO4, 45 mM NaCl, and 5 mM CaCl_2_, pH 7.8) on ColorFrost Plus slides (Fisher Scientific) coated with poly-L-lysine (Sigma Aldrich). Samples were freeze-cracked in liquid nitrogen. Sperm spreads to analyze detached spermatocytes and spermatids were obtained by applying slight pressure to the coverslip before freeze-cracking. Samples were fixed overnight in −20°C methanol. Specimen preparation and antibody labeling followed established protocols (Shakes et al., 2009). Primary antibodies included: 1:1250 rabbit anti-SPE-18; 1:600 4D5 mouse anti-MSP monoclonal (Kosinski et al., 2005), and 1:50 1CB4 monoclonal (Okamoto and Thomson, 1985). All samples were incubated with primary antibodies for 2 hours at room temperature. Affinity-purified secondary antibodies included 1:100 TRITC conjugated goat anti-rabbit IgG (Jackson ImmunoResearch Laboratories) and 1:100 FITC or DyLight-conjugated goat-anti-mouse IgG (H+L) (Jackson ImmunoResearch Laboratories). In some experiments, appropriately diluted working solutions of the secondary antibodies were preabsorbed with a powder made from acetone-fixed *C. elegans* (Miller and Shakes, 1995).

Final slides were mounted with DAPI containing Fluoro Gel II mounting medium (Electron Microscopy Sciences). Images were acquired under differential interference contrast or epifluorescence using an Olympus BX60 microscope equipped with a QImaging EXi Aqua CCD camera. Photos were taken, merged, and exported for analysis using the program iVision. For multi-dimensional imaging, z-axis stacks were taken using a z-axis stage controller at 0.2 mm intervals. For deconvolution, images were run through MicroTome deconvolution software. In some cases, the levels adjust function in Adobe Photoshop was used to spread the data containing regions of the image across the full range of tonalities.

For DIC/Hoechst preparations, males were dissected in buffer with 100 μg/ml Hoechst 33342 (Sigma Aldrich) on non-plus slides and immediately imaged.

### Western Blot

For western blot analysis, 100 worms were collected in 15-25 microliters of M9 buffer in the cap of a 1.5 microliter Eppendorf tube. Tubes were centrifuged for 1 minute at 15,000 rcf, immediately frozen in liquid nitrogen and stored at −80°C. Worm lysates from one freeze-thaw cycle were homogenized with a 4:100 mix of β-mercaptoethanol (MP Biomedicals) and sample buffer (NuPAGE LDS 4X Sample Buffer, Invitrogen) heated to 100°C, boiled for 5 minutes, and centrifuged for 8 minutes at 15,000 rcf. Lysates from 50-100 worms were loaded per lane, and proteins were resolved at 150V via SDS-PAGE (NuPage Novex 4-12% Bis-Tris, Invitrogen), and transferred to a PDVF membrane (GE Healthcare). After blocking overnight with pH 8.0 Tris-buffered saline with 0.1% Tween20 containing either 4% non-fat dry milk (Carnation) or 5% bovine serum albumin (Sigma-Aldrich), membranes were incubated with the appropriate primary antibody diluted in blocking buffer (4% milk or 5% BSA in 1X TBST) for two hours at room temperature, followed by incubation with 1:20000 peroxidase-conjugated secondary antibody (Abcam) for two hours at room temperature, and then developed by enhanced chemiluminescence (Immobilon Western Chemiluminescent HRP substrate, Millipore). SPE-18 protein was detected by a 1:5000 dilution of rabbit anti-SPE-18 polyclonal antibody (YenZym) and HRP conjugated goat-anti rabbit IgG (Abcam #ab6721). MSP was detected by a 1:10000 dilution of mouse anti-MSP monoclonal antibody 4A5 (Kosinski et al., 2005) and HRP conjugated goat anti-mouse IgG (#ab6789).

## ACKNOWLEDGEMENTS

Some strains were provided by the *Caenorhabditis* Genetics Center, which is funded by the NIH Office of Research Infrastructure Program [P40 OD010440]. We thank David Greenstein and Stephen L’Hernault for antibodies, and Harold Smith for the SPE-18 expressing *E. coli* strain. Lidia Epp (W&M molecular core technician) and undergraduates Alana Noritake and Bryan Neva assisted with the genetic and molecular identification of SPE-18 as F32A11.3. We thank Jordan Ward, Penny Sadler, and Kayleigh Morrison for critical reading of this manuscript.

## AUTHOR CONTRIBUTIONS

Conceptualization: D.C.S.; Methodology: D.C.S, K.L.P., M.S.P, and C.M.U.; Validation: K.L.P., M.S.P, and C.M.U; Formal analysis: D.C.S, K.L.P., M.S.P, and C.M.U.; Investigation: K.L.P., M.S.P, and C.M.U.; Writing - original draft: M.S.P., K.L.P., D.C.S.; Writing - review & editing: K.L.P., M.S.P., D.C.S.; Visualization: K.L.P., D.C.S.; Supervision: D.C.S.; Project administration: D.C.S.; Funding acquisition: D.C.S.

## DATA AVAILABILITY

Not applicable

